# Phage co-transport with hyphal-riding bacteria fuels bacterial invasion in water-unsaturated microbial ecosystems

**DOI:** 10.1101/2021.07.28.454208

**Authors:** Xin You, René Kallies, Ingolf Kühn, Matthias Schmidt, Hauke Harms, Antonis Chatzinotas, Lukas Y. Wick

## Abstract

Non-motile microbes enter new habitats often by co-transport with motile microorganisms. Here, we report on the ability of hyphal-riding bacteria to co-transport lytic phages and utilize them as ‘weapons’ during colonization of new water-unsaturated habitats. This is comparable to the concept of biological invasions in macroecology. In analogy to invasion frameworks in plant and animal ecology, we tailored spatially organized, water-unsaturated model microcosms using hyphae of *Pythium ultimum* as invasion paths and flagellated soil-bacterium *Pseudomonas putida* KT2440 as carrier for co-transport of *Escherichia* virus T4. *P. putida* KT2440 efficiently dispersed along *P. ultimum* to new habitats and dispatched T4 phages across air gaps transporting ≈ 0.6 phages bacteria^−1^. No T4 displacement along hyphae was observed in the absence of carrier bacteria. If *E. coli* occupied the new habitat, T4 co-transport fueled the fitness of invading *P. putida* KT2440, while the absence of phage co-transport led to poor colonization followed by extinction. Our data emphasize the importance of hyphal transport of bacteria and associated phages in regulating fitness and composition of microbial populations in water-unsaturated systems. As such co-transport mirrors macroecological invasion processes, we recommend hyphosphere systems with motile bacteria and co-transported phages as models for testing hypotheses in invasion ecology.

## Introduction

To cope with the heterogeneous and highly changeable soil environment, microbes have evolved inter-microbial co-transport strategies to gain motility and colonize new habitats (reviewed by[1]). For instance, bacteria have been found to efficiently disperse along hyphae in soil. Fungi embody up to 75% of the subsurface microbial biomass. Their hyphae create fractal mycelial networks of 10^2^ to 10^4^ m per g of topsoil and efficiently explore heterogeneous air-filled habitats[2–4]. They thereby serve as pathways for bacteria to efficiently disperse (fungal highways[5, 6]), forage[7] and colonize new habitats [8, 9]. Hyphae however may reduce dispersal of intrinsically non-motile yet abundant (>10^8^ g^−1^ soil[10]) soil virus-like particles as they have been shown to retain waterborne phages [11, 12]. Considering the slow diffusion (~ 0.034 mm/d[13]) and enhanced inactivation at dry conditions[14, 15], transport of phages seems particularly restricted in water-unsaturated habitats. Recent studies have revealed that phages in aquatic environments can adsorb to surfaces[16, 17], mucus[18], flagella[19] of non-host bacteria or even sheath surrounding them[20]. As hyphae are adapted to water-unsaturated habitats we here hypothesize that hyphae allow for phage co-transport with hyphal-riding bacteria and thereby fuel the fitness of invading bacteria in new regions (alien ranges) of water-unsaturated habitats. Co-transported phages, which are not specific to the carrier bacteria but specific to resident bacteria in alien ranges thereby may serve as “biological weapons”[21, 22] and increase the competitive ability and fitness of the invading carrier bacteria. Temperate phages integrated in bacterial genome (i.e. as prophages) have been suggested to serve as agents of ‘bacterial warfare’ [22–24]. In aquatic environments, lytic phages adsorbed to bacteria have been shown to facilitate phage infection of biofilm bacteria and to promote biofilm colonization by carrier bacteria [19]. In unsaturated environments like soil however, little is known about co-transport of phage with motile bacteria and its effect on bacterial population dynamics.

In analogy to the previously published MAcroecological Framework of Invasive Aliens (MAFIA[25]) we here tailored spatially organized microcosm systems to mimic the stages of the invasion process (i.e. transport, introduction, establishment, and spread) [25–28] of co-transported phages and bacteria in the hyphosphere. Unlike biological invasions in macroecological systems, our model system bears the advantage that all interacting species, their locations and location characteristics as well as invasion events are known and can be manipulated. In our model the ‘native range’ and the ‘invaded’ or ‘alien range’ are two agar patches that are separated by an air gap. The air gap serves as a barrier that only can be overcome by bacterial movement along hyphae as invasion pathways crossing the ‘native’ and ‘alien’ ranges. Such water-unsaturated systems allow (i) to evaluate the transport efficiency of flagellated and non-flagellated bacteria as vectors to transport phages into alien ranges, and (ii) to quantify possible population effects of co-transported phages in the alien range. Our approach revealed that motile bacteria can help phages to migrate into water-unsaturated habitats and that phage co-transport can fuel the settlement and fitness of hyphal-riding bacteria in host pre-colonized alien ranges.

## Materials and methods

### Strains, growth conditions and enumeration methods

GFP-labeled wild type [29] soil bacterium *Pseudomonas putida* KT2440 (termed hereafter as WT) and its non-flagellated mutant Δ*filM* were used as carriers for phage co-transport. Δ*filM* was obtained by allelic exchange with a truncated version of *filM* [30] and used to test the role of flagella for phage sorption and phage co-transport. Both strains were kindly provided by Arnaud Dechesne (DTU). They were cultivated in LB medium on a gyratory shaker at 30°C and 150 rpm. For microcosm experiments, an overnight culture (OD600 ≈ 2) was washed once with PBS buffer (100 mM) and adjusted to reach an OD600 ≈ 4 (≈ 8 x 10^6^ cell μL^−1^). Hyphae of the oomycete *Pythium ultimum* [31] were used as model dispersal networks. It was pre-grown on potato dextrose agar (PDA) at room temperature (RT) [32]. *Escherichia* virus T4 (T4) was selected as phage in co-transport experiments. T4 was propagated on its host *E. coli* (Migula 1895) using the liquid broth method in DSM544 medium [33]. T4 and *E. coli* were purchased from Deutsche Sammlung von Mikroorganismen und Zellkulturen GmbH (DSMZ, Braunschweig, Germany). *E. coli* was cultivated in DSM544 medium at RT on gyratory shaker at 150 rpm (generation time = 41 ± 0.1 min in the exponential phase). For microcosm experiments, 10 μL of an overnight *E. coli* culture were transferred into 20 mL fresh DSM544 medium and cultivated at 30°C until early exponential growth (OD_600_ ≈ 0.4).

Enumeration of *E. coli* and *P. putida* KT2440 were carried out by counting colony forming units (CFU) on LB agar incubated at 30°C overnight. When both strains were present in a sample, CFU of GFP-tagged *P. putida* KT2440 were counted with an epifluorescence microscope equipped with a black-and-white camera (AZ 100 Multizoom; Nikon, Amsterdam, Netherlands) under the GFP channel using NIS Elements software. Plaque forming unit (PFU) enumeration was done using a modified small-drop plaque assay technique as detailed earlier [34] allowing the double-layer counting plates to be incubated overnight at 37°C. The whole-plate plaque assay (cf. [35]) was also performed to crosscheck PFU counts for samples with zero PFU count by small-drop plaque assay.

### Determination of phage adsorption to bacteria

Adsorption efficiencies of T4 to WT, Δ*filM* and *E. coli* were quantified at phage-to-bacteria ratios of 1, 0.1 and 0.01 in 6-8 replicates as described earlier [36]. In brief, suspensions of bacteria (≈ 10^8^ CFU mL^−1^) and T4 were incubated in PBS at RT for 1 h (15 min for T4 and *E. coli*) and centrifuged at 8,000 × g at 4°C to pellet bacteria and adsorbed phages. Amounts of adsorbed phages were estimated by the loss of free phages after centrifugation; i.e. phage adsorption (%) calculated by the ratio of adsorbed phages to total phages prior to centrifugation. A phage-only control was also included to determine the stability and change of infectivity of T4 in the medium and during centrifugation.

### Quantification of phage co-transport by hyphal-riding bacterial carriers

T4 co-transport with carrier bacteria was quantified in quintuplicate laboratory microcosms mimicking water-unsaturated soil in the presence and absence of hyphal networks (Fig 1a). The microcosms consisted of an agar patch A (PDA, 2% agar (w/v), l × w × h = 1 × 1 × 0.6 cm) that was separated from an agar strip (l × w × h = 2 × 1 × 0.6 cm) by a 0.5 cm air gap. *P. ultimum* was pre-inoculated on agar patch A for 3-4 days to reach > 0.5 cm hyphal length prior to finally assembling the microcosms. The agar strip (that was split into equally sized patches B & C before harvesting, Fig. 2a & 3a) however was freshly prepared upon setting up the microcosm. It was made from minimal medium agar (MMA) to avoid bacterial growth and consisted of a top layer (MMA, 0.6% agar (w/v), h = 0.1 cm) and a bottom layer (MMA, 2% agar (w/v), h = 0.5 cm). All agar patches were placed in sterile Petri dishes. To analyze the transport of T4 along hyphae of *P. ultimum* in presence and absence of carrier bacteria, five different scenarios were used (Fig. 1b): (i) WT, (ii) WT + T4, (iii) T4, (iv) Δ*filM*, and (v) Δ*filM* + T4. *P. ultimum* pre-grown agar patches (equal size as agar patch A) with T4 or WT + T4 were used to quantify hyphal effects on T4 infectivity. Inactivation of T4 on agar surfaces was studied on agar patches in the absence of *P. ultimum*. In scenarios (ii) and (v) bacteria with previously adsorbed phages were added. To do so, T4 (6 × 10^9^ PFU mL^−1^) was co-incubated in PBS with WT or Δ*filM* (≈ 8 × 10^9^ cells mL^−1^) at RT for 1 h at 125 rpm and then centrifuged (8,000 × g for 10 min at 4°C) to discard free phages in the supernatant. The remaining pellet containing bacteria and adsorbed phages was washed once and concentrated by re-suspension in PBS to reach an inoculum density of OD_600_ (estimation) ≈ 20. Inocula with either bacterial cells (≈ 4×10^10^ cells mL^−1^) or phages (≈ 6×10^8^ PFU mL^−1^) in PBS served as controls. 1 μL of the respective inoculum (i.e. ≈ 4×10^7^ bacteria or 6×10^5^ phages) was placed at 0.25 cm from the left edge of agar patch A. After inoculation the Petri dishes were sealed with Parafilm®, placed in a plastic container and incubated at 20°C in the dark. After 24, 48 and 72 h the microcosms were sacrificed and phage and bacteria numbers quantified on agar patches A, B & C by PFU and CFU. Isolation of phage and bacteria from agar was done as described previously [7]; i.e. cut agar pieces were suspended in 3 mL PBS in glass tubes, vortexed at maximal speed for 1 min and then sonicated (2 × 30 s with a break of 1 min). Phage-bacteria suspensions were 1:1 extracted with chloroform in order to inactivate and remove bacterial biomass prior to PFU quantification.

**Figure 1.**
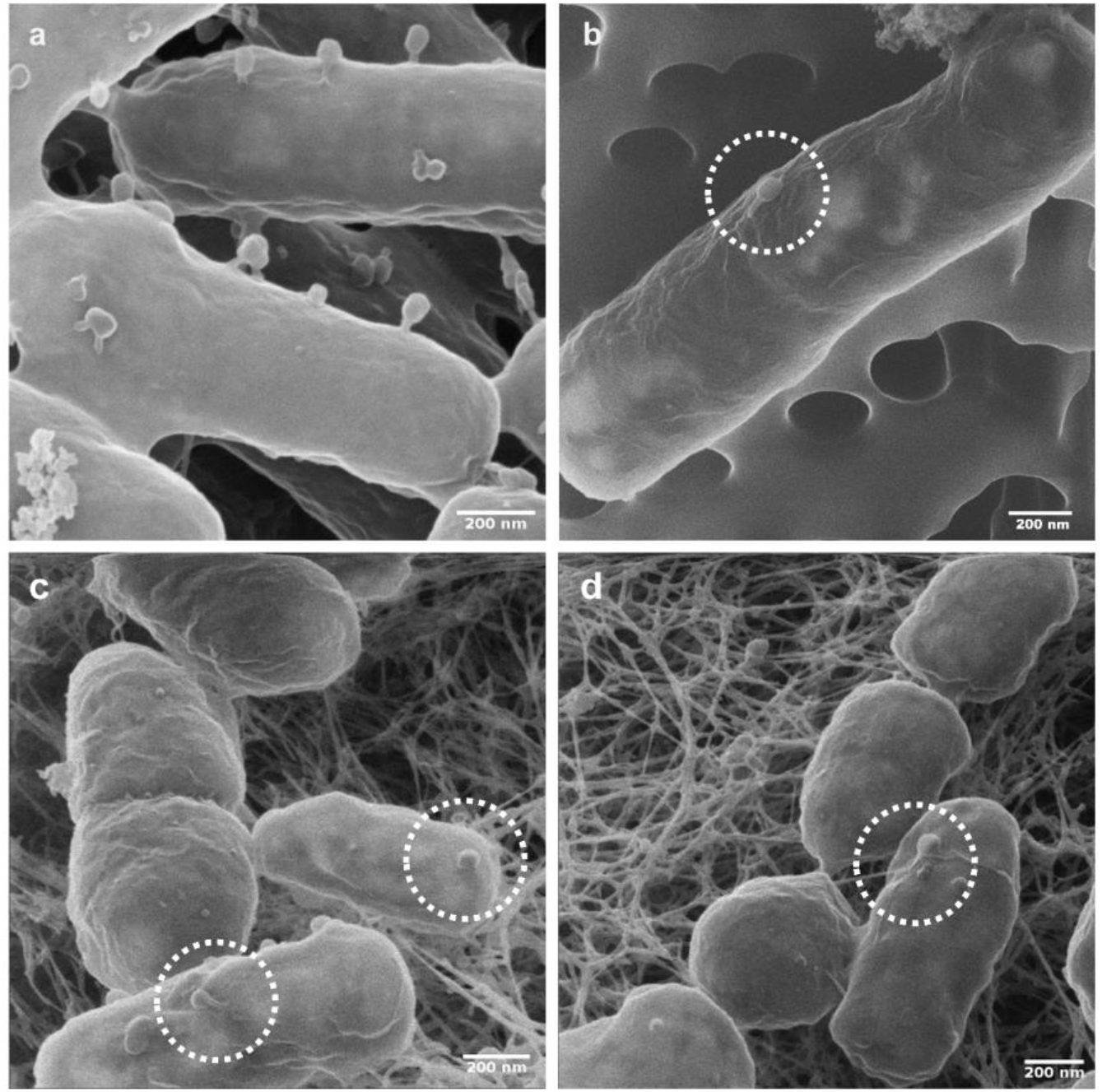
Helium ion microscopy (HIM) visualization of T4 phage adsorption to *E. coli* host and non-host *P. putida* KT2440 cells. **Figs 1a&b** visualize cells from phage adsorption experiments (cf. materials and methods) reflecting tail-mediated adsorption to *E. coli* host cells (**Fig. 1a**) and capsid-driven adsorption to non-host *P. putida* KT2440 (**Fig 1b**). **Figs 1c&d** visualize tail- (**Fig-1c**) and capsid-driven (**Fig. 1d**) phage adsorption to biofilm cells growing on agar patch B on day 2 in experiments evaluating population effect of T4 co-transport with *P. putida* KT2440 (cf. Fig. 3).

### Evaluation of population effects due to phage co-transport with hyphal riding bacteria

Effects of phage co-transport on the populations of carrier bacteria invading an alien range occupied by competing bacteria were evaluated in five replicates in similar microcosms as described above. The double-layer of agar patches B & C, however, contained nutrient agar (DSM544) and the upper agar layer was densely populated by *E. coli* (Fig. 3a). Prior to the experiment, the thin upper agar layer of agar patches B & C was inoculated with *E. coli* (5 ± 0.5 × 10^4^ CFU cm^−2^) and allowed to grow at RT for 3 h (≈ 4 generations, 2.3 ± 0.1 × 10^5^ CFU cm^−2^). Three different scenarios were studied using either (i) T4, (ii) WT, or (iii) WT +T4 as inocula. After 24, 48, and 72 h, all agar patches A, B & C were harvested and phages and bacteria quantified as described above.

### Helium Ion Microscope Imaging

Helium Ion Microscope (HIM) imaging was performed with suspensions from phage-bacteria adsorption assays and surfaces of agar patches B after invasion of phage-carrying WT *P. putida* KT2440 into *E. coli* populations. HIM imaging of surfaces of T4 plaques on a lawn of *E. coli* served as control. Suspension containing phage-bacteria associations from the adsorption assays were mixed at a ratio of 1:1 with 2% (v/v) paraformaldehyde in 0.2 M sodium‐cacodylate buffer (pH = 7.4) and allowed to stand for 2 h for chemical fixation. The suspension was then filtrated onto 0.22-μm pore‐size polycarbonate filter papers (Merck‐Millipore) using a Sartorius hand‐filtration device. The filter papers were rinsed twice for 5 min with Na‐cacodylate buffer to remove salts and debris. The samples were dehydrated in a graded aqueous ethanol series (30%, 50%, 70%, 80%, 90%, and 100% EtOH) and critical point dried. To observe phage-bacteria associations on agar surfaces, the agar patches were submersed in Petri-dishes in the fixative for 2 h (cf. above). The fixative was then gradually exchanged using a graded aqueous ethanol series (>50% fixative replacement each time to avoid material loss) and the sample finally critical point dried. The dried samples were mounted onto standard stubs for electron microscopy using a conductive silver epoxy glue and imaged by a Zeiss Orion NanoFab (Zeiss, Peabody, MA, USA) scanning helium ion microscope using an ion-landing-energy of 25 keV, a 10-μm aperture and an Everhard-Thornley-type secondary electron detector. To achieve both high lateral resolution (≤ 2 nm) and contrast, the beam current was set between 0.08 pA (high magnification) and 0.25 pA. Charge compensation during imaging was achieved with an electron flood-gun operated in line-flooding mode. In order to avoid beam damage and to allow for efficient charge compensation the dwell time of the beam on a pixel was kept between 0.5 −1.0 μs.

### Data analysis and statistics

The time-dependent transport rate (*R*_i_; shown in eq.1) of phages (*R*_p_, PFU cm^−1^ d^−1^) or bacteria (*R*_b_, CFU cm^−1^ d^−1^) was obtained by normalizing the number of phages transported (*N*_p_, PFU) or bacteria transported (*N*_b_, CFU) to the dispersal distance (*d*, cm) and the time (*t*, d) until harvesting. Because T4 got rapidly inactivated on agar surfaces (99% loss of PFU in < 24 h, Fig. S2a) and subsequent low phage numbers were elusive to direct quantification, *N*_p_ was approximated by the difference of phages counted at the point of inoculation (i.e. agar patch A) in the absence and presence of carrier bacteria. This approach was possible as the presence of either *P. ultimum* or *P. ultimum* and *P. putida* KT2440 co-cultures did not influence T4 infectivity and enumeration (Fig. S2c).

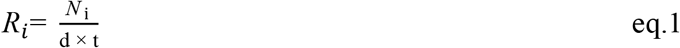

For co-transported phages, time-dependent transport capacity (*C*_p_, PFU bacteria^−1^) and transport efficiency (*E*_p_, %). reflects the average number of phages transported by a single bacterium and *E*_p_ the fraction of phages dispatched by carrier bacteria. For calculation details, please refer to extended materials and methods in SI.

To evaluate the effect of phage co-transport on bacterial or phage populations at time *t*, the absolute fitness (*W*_i_)[37] of a given population i was calculated using eq. 2. *W*_i_ is the time dependent ratio of the population size (bacteria or phages) on given agar patches in the presence of *E. coli* (*N*\*_i_, CFU or PFU) and absence of *E. coli* (*N*_i_,, CFU or PFU). *W* > 1 and *W* < 1 indicate an increase and a decrease of the population size, while *W* = 0 refers to population extinction (c.f. extended materials and methods in SI for calculation details).

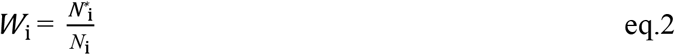

Data were plotted as transparent dots and statistics were displayed as median (circle) with 95% confidential interval (95CI, vertical bar) or boxplot notches as modified from [38]. The 95CI was determined by bootstrapping (1000 samples) and derived from the 2.5^th^ and 97.5^th^ percentile[39]. When the 95CIs of two conditions do not overlap, it indicates a statistical difference between these two conditions[40].

## Results

### Adsorption of T4 phage to WT and ΔfilM

Being a prerequisite for phage co-transport with carrier bacteria we tested the adsorption of T4 phages to the flagellated WT of *P. putida* KT2440 and its non-flagellated Δ*filM* mutant. We found that 13 - 62% of T4 particles adsorbed to the non-host WT with the highest adsorption (62%) observed at a phage to bacteria ratio of 1 (Fig. S1a). No statistically significant differences (P > 0.05) between T4 adsorption to flagellated WT and Δ*filM* were observed at this phage to bacteria ratio (Fig. S1b). Adsorption to WT was thus consistently lower and more variable than to the host strain *E. coli* (68 - 84%; Fig. S1a). T4 adsorption was further evidenced by HIM visualization. It revealed capsid-driven adsorption of T4 (Figs. S1d&e) to the surface of *P. putida* KT2440 leaving the phages’ tails unattached (Fig. 1b). This is in contrast to T4 adsorption to host *E. coli*, where perpendicular adsorption with phage tails bound to bacterial surfaces was found (Fig. 1 & Fig. S1c).

### Effect of hyphal-riding bacteria on phage co-transport

In analogy to the different stages of the plant or animal invasion processes [25, 26] we developed a spatially organized microcosm system to evaluate phage co-transport with hyphal-riding bacteria invading new habitats (i.e. agar patches B & C; Fig. 2a). To challenge the role of bacterial motility for phage co-transport, we used flagellated WT and the non-flagellated Δ*filM* mutant in five different scenarios (Fig. 2b): (i) T4, (ii) Δ*filM*, (iii) WT, (iv) Δ*filM* + T4, and (v) WT + T4. No airborne transport of phages was observed in the microcosms (Fig S2b). As T4 infectivity got rapidly lost on agar surfaces (> 99% loss within 24 h, Fig. S2a) preventing reliable enumeration on agar patches B & C, phage transport rates (*R*_p_; eq. 1) were calculated by differences of phage counts on agar patch A in all scenarios. Contrary to our observation on sterile agar surfaces (Fig. S2a) T4 maintained its infectivity up to two days when placed on agar patches covered by *P. ultimum* (Fig 1c). No reduction in T4 counts in the absence of bacteria or in presence of Δ*filM* was observed over two days pointing at negligible diffusion of viable phages or phage co-transport by Δ*filM*. Similarly, inoculation of WT + T4 on isolated *P. ultimun* agar (i.e. no WT exportation; Fig S2c) showed no reduction in T4 counts (Fig. S2c). By contrast, inoculation of WT + T4 on interlinked *P. ultimun* agar (i.e. WT exportation allowed; Fig. 2a) resulted in a significant (> 65%) reduction of T4 counts after two days and a transport efficiency of *E*_p_ ≈ 60% (Fig. 2e). The phage transport rates (*R*_p_ ≈ 10^5^ PFU cm^−1^ d^−1^, Fig. 1d) thereby coincided with hyphal dispersal rates of the WT (*R*_b_ ≈ 1.4 × 10^5^ CFU cm^−1^ d^−1^; Fig. 1d) suggesting an apparent transport capacity of *C*_p_ ≈ 0.6 PFU bacteria ^−1^ after 2 days (Fig. 1g). After three days a 10-fold decreased T4 transport rate (*R*_p_ ≈ 1.4 × 10^4^ PFU cm^−1^ d^−1^, Fig. 1d) and a 20-fold reduced apparent T4 transport capacity (*C*_p_ ≈ 0.03 PFU bacteria^−1^) were observed. This is likely due to growth of WT bacteria and/or the inability of the WT progeny on agar patch A to get into contact with T4 phages. Transport rates of WT bacteria were similar regardless of the presence of T4 (Fig. 2e).

**Figure 2.**
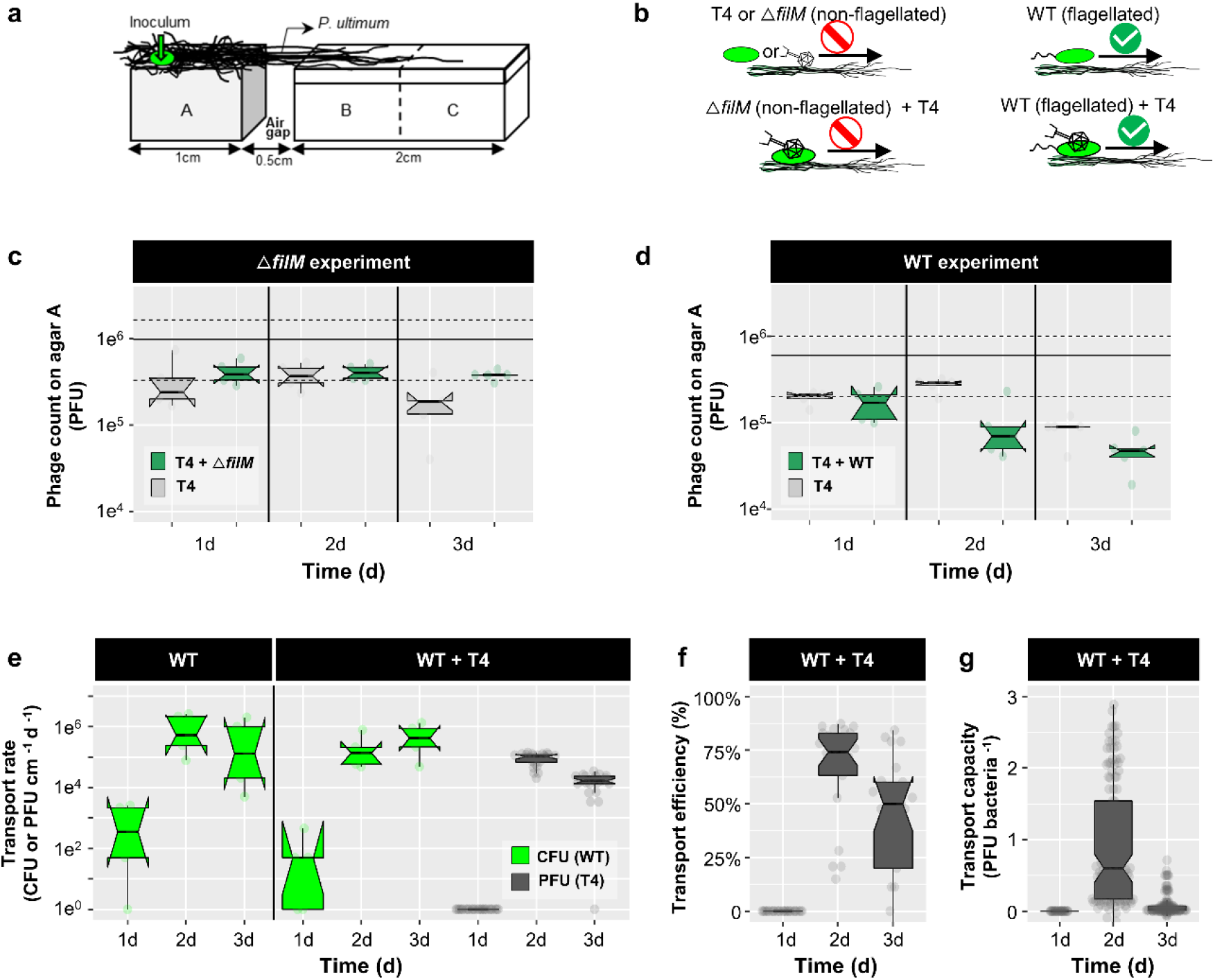
Co-transport of T4 with *P. putida* KT2440 (WT or non-flagellated Δ*filM*) along hyphae of *P. ultimum*. **Fig 2a.** Scheme of the microcosm setup specifying microbial inoculation points and the spatial arrangement and dimensions of agar patches A, B & C. **Fig 2b**. Experimental scenarios and observed results. Upper left panel: T4 or Δ*filM* do not disperse along hyphae over the air-gap. Lower left panel: Δ*filM* does not transport T4 along hyphae. Upper right panel: WT disperse along hyphae. Lower right panel: WT disperses along hyphae and transports T4 along hyphae over the air-gap. **Figs 2c&d**. T4 counts on agar patch A after one day in the absence and presence of Δ*filM* (**Fig. 2c**) and WT (**Fig. 2d**). The solid and dashed lines indicate the median of 5 replicates and its 95CI of the initial inocula. **Fig 2e**. Time dependent cumulative transport rates of WT (R_WT_, in green) and phages (*R*_T4_, in grey) to agar patches B&C (cf. eq. 1). **Fig 2f.** Time dependent phage transport efficiency in presence of WT (*E*_T4_). **Fig 2g.** Time dependent phage transport capacity of WT (*C*_T4_). Data on T4 transport by Δ*filM* are not shown as no transport was observed. Notches of the boxes represent 95CI of 5 replicates. If notches of two conditions do not overlap, it indicates a statistical difference between the two conditions.

### Effects of phage co-transport on bacterial invasion and invader fitness

To evaluate the community effect of T4 co-transport with hyphal-riding WT, agar patches B & C were covered with 2.3 ± 0.1 × 10^5^ CFU cm^−2^ of *E. coli* as local host bacteria of T4 (Fig. 3a). Development of T4, WT and *E. coli* counts was quantified over time in four different scenarios (cf. Fig. 3b): (i) PBS only, (ii) WT, (iii) T4, and (iv) T4 + WT. In the absence of WT, no diffusion of infectious T4 along hyphae to agar patches B & C was observed at any time (Fig. 3c). In presence of WT, however, ~10^5^ PFU and ~10^4^ PFU were recovered from agar patches B & C after 1 day leading to a 45-180-fold increased phage abundance (Fig. 3c; 45< *W*_T4_ <180 as defined by eq. 2). The presence of T4 went along with HIM-detectable bacterial lysis of *E. coli* (Fig. S3e) indicating previous phage lysis on day 1. No T4, however, were detected on day two (Fig. 3c) despite HIM-detectable phages adsorbing to bacterial surfaces either in tail-mediated perpendicular (Fig. 1c) or capsid-mediated coaxial positions (Fig. 1d). Tail-mediated adsorption thereby points at an infection of host cells (Fig. 1a) while capsid-mediated adsorption may refer to unspecific interaction with non-host cells (Fig. 1b). In the absence of T4 the carrier WT cells invaded and established on agar patches B & C in the first 24 h (*W*_WT_ > 930) accounting for 0.08-0.39% of all bacteria. Thereafter WT got strongly inhibited (*W*_WT_ < 0.06) and comprised less than 0.01% of the population; this, despite a constantly growing WT population on agar patch A (*W*_WT_ > 1) and a likely on-going WT invasion from agar patch A (Fig 3c).

**Figure 3.**
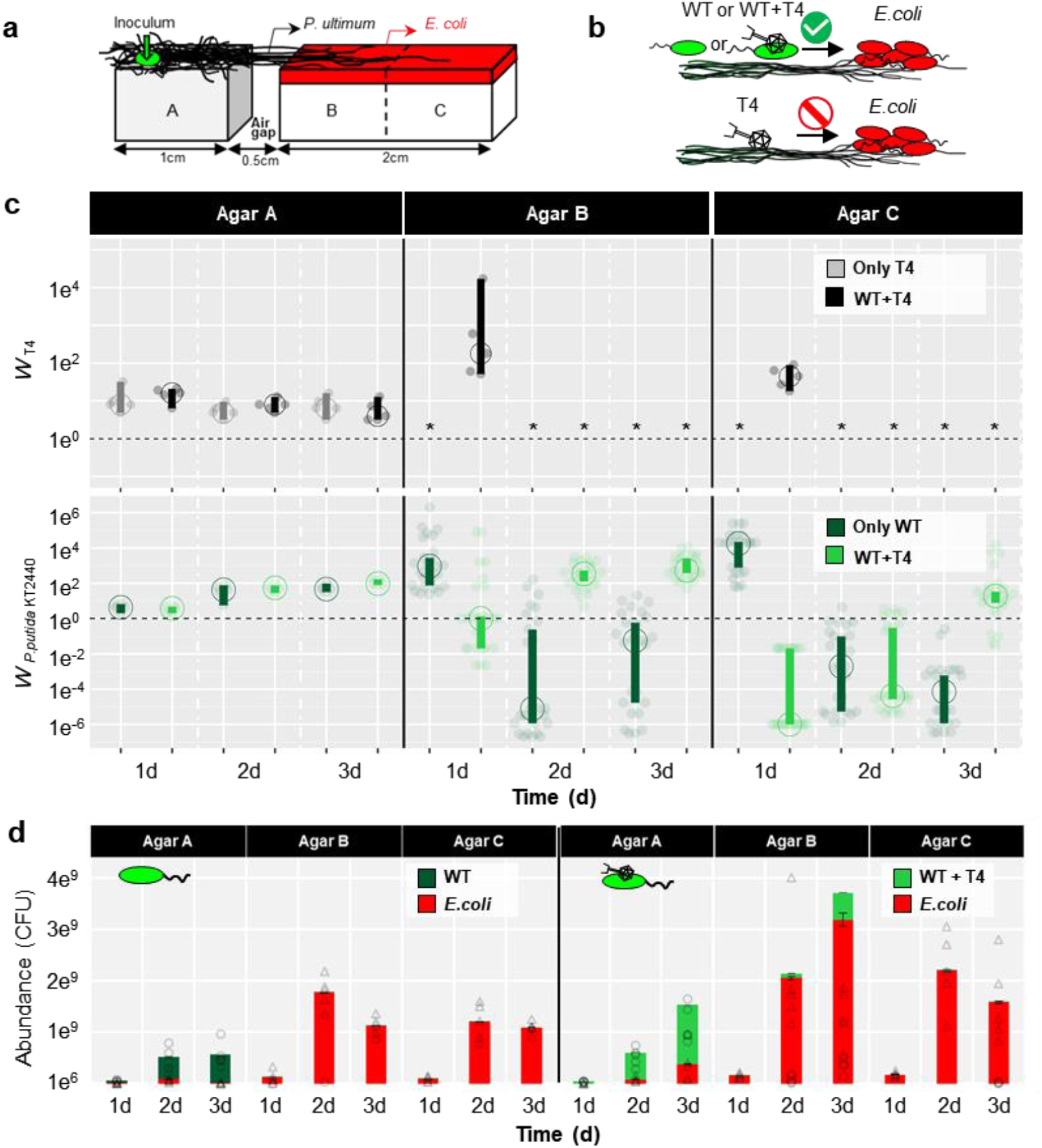
Effects of T4 co-transport with flagellated *P. putida* KT2440 (WT) along hyphae of *P. ultimum* phage, *E. coli* and WT counts on agar patches A-C. **Fig 3a.** Scheme of the microcosm setup specifying microbial inoculation points and pre-colonized *E. coli* on agar patches A, B & C. **Fig 3b.** Overview of the scenarios for evaluating invasion. Upper panel: T4 co-transports with WT along hyphae to an *E. coli* covered alien range (i.e. agar patches B & C). WT disperses along hyphae to the *E. coli* pre-colonized alien range. Lower panel: T4 is not able to disperse along hyphae to the alien range. **Fig 3c.** Time dependent absolute fitness of T4 (*W*_T4_) and of *P. putida* KT2440 (*W*_WT_) on agar patches A, B & C in presence and absence of phage-bacterial co-transport. Dashed line represents no change of the population size (*W*_i_ = 1) and * indicates that no PFU were detected. The vertical bars represent 95CI calculated from 5 biological replicates. If vertical bars of two conditions do not overlap, it indicates a statistical difference between the two conditions. **Fig 3d.** Abundance of *E.coli* and WT populations on agar patches A, B & C with and without T4 co-transport (triangles and circles represented individual data points of *E.coli* and *P. putida* KT2440 in 5 replicates). Please note that abundances of WT < 10^8^ CFU (e.g. observed in the case of WT + T4” on 3d) are not visible due to the plot scale.

By contrast, T4 co-transport with WT promoted invasion and fitness of the carrier bacteria after a delay of one day: on agar patch B the absolute fitness of WT increased from *W*_WT_ ⩽ 1 (1 d) to *W*_WT_ = 556 (≈ 5 × 10^8^ CFU) at t = 3 d (Fig 3c). On agar patch C the WT fitness changed from *W*_WT_ < 1 (t = 1 −2 d) to *W*_WT_ > 1 after 3 days. After 3 days WT accounted for ≈ 14% and 1% of the total bacterial population on agar patches B and C, resp. (Fig. 3d). Epifluorescence microscopy analysis thereby revealed that gfp-labeled WT established best in the vicinity of the hyphae (Figs. S3b & c), suggesting highest phage-clearance effects on resident *E. coli* in the hyphosphere. Invasion of WT, however, had no negative impact on the abundances of *E. coli* cells in agar patches B & C (Fig. 3d). We further checked the effect of phage co-transport on overall phage counts and abundances of host and carrier bacteria in the microcosms (i.e. agar patches A, B & C) (Fig. S4). While co-transport did not influence T4 abundances (Fig. S4a), it clearly enhanced the abundance of WT (Fig. S4b), yet not influence the *E. coli* abundances (Fig S4c).

## Discussion

### Phage co-transport with hyphal-riding bacteria

Although described for aquatic environments [19, 41, 42], little is known on viral co-transport with non-host microorganisms in vadose habitats; even though adsorption and subsurface co-transport of nano-colloids with organic or biological materials has been described nearly two decades ago [43]. Here, we show that *Escherichia* virus T4 can adsorb to the hyphal-riding soil bacterium *P. putida* KT2440 (Fig. S1). It thereby gets efficiently dispersed in water-unsaturated environments (Fig. 2) and invades new habitats (Fig. 2a) even if this involves crossing air-filled spaces (Fig 1c). No diffusion of T4 along hyphae of *P. ultimum* and no T4 co-transport with the non-flagellated Δ*filM* was detected. Flagellar mobility hence seems to be a driver for efficient phage transport by bacteria. Although a few studies have reported on preferential adsorption of phages to bacterial flagella [19], we found similar efficiencies of T4 adsorption to flagellated WT and non-flagellated Δ*filM* (Fig S1). T4 adsorption to the bacterial cell surface rather than to flagella was also confirmed by HIM imaging. Microscopy suggested that capsid-mediated sorption to cells left the phage tails unattached. This allows adsorbed phages to remain infectious, i.e. transferable to thermodynamically favored adsorption to host cells [44]. We chose T4 as model phage as it is known to adsorb to cell surfaces and to be less adsorptive than smaller phages [45, 46]. T4 adsorption to and co-transport by carrier bacteria hence may therefore lead to conservative estimates for phage adsorption and co-transport. For instance, a recent study using *Bacillus cereus* as a model bacterium for waste water has shown its ability to adsorb phages of different morphologies from 10 viral families [19]. T4, furthermore, was also found to adsorb to the soil bacterium *P. fluorescence* Lp6a despite of its highly distinct physico-chemical surface properties [47] from *P. putida* KT2440 (Fig S1a). As hyphal-riding bacteria are widespread in many environments (e.g. soil [48–50], the vicinity of roots [6, 51] and cheese[52]), we hence speculate that other combinations of phages with hyphal-riding carrier bacteria may also lead to phage co-transport. As phage co-transport with non-host bacteria can take place over days (Fig. 2) one may speculate that such co-transport may be more efficient than recently reported co-transport of phages with their host cells (‘virocells’) during the (typically small, e.g. ≥ 20 min for T4) latent period [53].

### Benefits of phage co-transport for bacterial carriers

Our data show that T4 co-transport fosters fitness of hyphal-riding WT cells if invading alien ranges occupied by resident T4 host cells (Figs. 3 & 4). Over 48 h the invasion and colonization of WT occurred in close vicinity (i.e. in the hyphosphere) of colonizing hyphae acting as preferential transport pathways for bacteria carrying adsorbed lytic phages (Fig. 4). Co-transported phages provided significant fitness gain for carrier bacteria in the alien range after one day (Fig. 3c). Yet, no phages could be detected at later stages although HIM analyses clearly revealed both phage particles adsorbing to bacterial surfaces (Figs. 1c & d) and lysed *E. coli* cells (Fig. S3e) in the hyphosphere (Fig. S3b). Such increase of the T4 abundance during initial WT carrier invasion may have been promoted by high accessibility of *E.coli* cells to T4 introduced by hyphal-riding WT. Lysis of host cells coupled with exponential growth of carrier WT cells in the hyphosphere however may have reduced *E. coli* cell density, and, hence, T4 accessibility and subsequent host infection. Similar phenomenon has been reported in spatially organized biofilms with two *Pseudomonas* strains, where the growth of the phage insensitive strain largely reduced the phage abundance by blocking their access to their host [54]. Likewise, on-going T4 transport may have triggered anti-phage mechanisms in the *E. coli* biofilms [55, 56]. Our data indicate that initial WT invasion in the absence of T4 was more efficient (*W*_WT_ > 930) than in its presence (*W*_WT_ ≈ 1). As transport rates of WT and WT + T4 did not differ (Fig. 2e, P > 0.05), we speculate that phage infection may lead to growth inhibition of ‘bystander bacteria’ as demonstrated in *Enterococcus faecalis* [57]. The hyphal-riding invader WT cells however could not establish in the absence of T4 - likely due to higher fitness of resident *E. coli* – and got eliminated after initial colonization success (Figs. 3c&d; Fig. 4).

**Figure 4.**
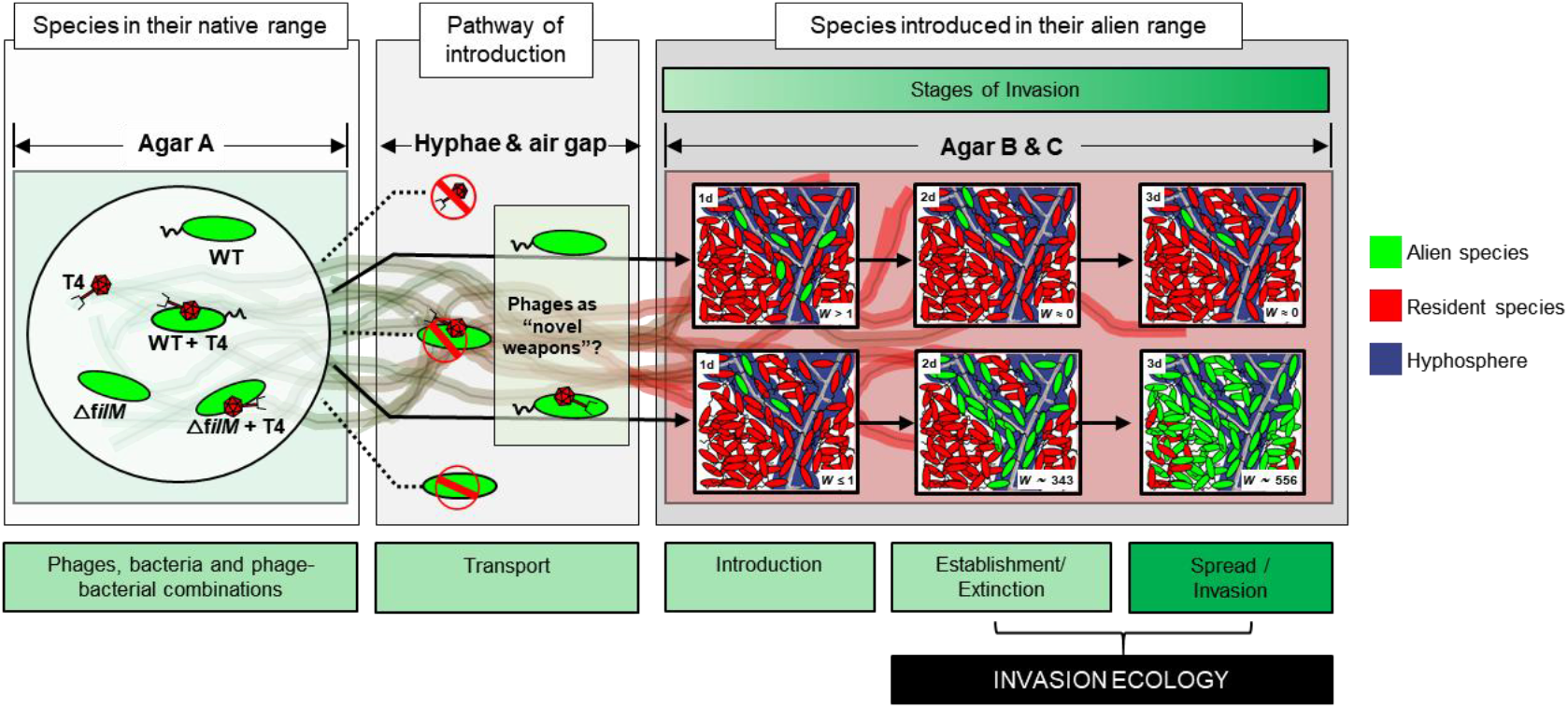
Equivalence between the MAcroecological Framework of Invasive Aliens (MAFIA) and the microsphere model system used in this study (cf. Fig. 3a). Based on the background sketch of the microsphere model system, the main findings of the study are summarized and illustrated in the flow chart. Boxes and contents are directly mirrored from the recently published MAcroecological Framework of Invasive Aliens[25] built upon [26–28].

### Microbial co-transport as a model for studying biological invasion

In analogy to the MAcroecological Framework of Biological Invasions [25, 26], we tailored a microbial system to study transport, introduction, establishment and spread of hyphal-riding bacteria in presence and absence of co-transported phages (Fig 4). The hyphosphere of fungi and oomycetes has been described to serve as pathway and scaffold for microbial transport (‘logistics’[9]) and evolution[58]. The role of ‘hitchhiker’ [1] phages as ‘weapon-like’ [21] promoters of microbial invasions and population dynamics in vadose habitats has not been described. Our findings hence may resemble the concept of “novel weapon” (e.g. biochemical possessed by invading species that is fatal to resident species) in plant ecology [21] or reflect the spread of infectious disease in animal ecology (e.g. Grey squirrely being vectors for squirrel pox infecting European red squirrels [59]). Such mechanisms have been difficult to be directly demonstrated in ecology, mostly due to limited data on population dynamics or inadequate observation time crossing multiple generations, especially with the respect to wild life populations[60]. By adapting spatially organized microbial model systems, those difficulties can be easily solved: multiple fast and reliable quantification methods, ranging from culture-based enumeration (this study) to molecular methods (e.g. qPCR) or –omics approaches, can be employed to resolve the microbial population dynamics. Hundreds of generations (e.g. > 10^4^ generation of *E. coli* within 3 days) can be covered in a few days and spatial organization can be overseen by microscopic imaging. In our study for example we evidenced the effects of phage-bacteria co-transport on invasion success in experiments as short as three days. In such a system, we can easily manipulate not only the stages of the invasion process e.g., transport/pathway (transport rates *R*_i_, capacity *C*_i_, efficiency *E*_i_) or introduction [25–28], by changing the barrier to be overcome, as well as the medium (e.g., hyphae) to cross this barrier. We can also manipulate species traits (by choosing bacteria and/or phages with different traits), location characteristics (e.g., changing nutrient content as well as adding non-beneficial substances such as poisons), event characteristics (e.g., propagule pressure by changing population sizes *N*) and any of their interactions affecting fitness *W*. As phage interaction with non-host bacteria seems to be a widespread mechanism[19], future exploration of hyphae-associated phage-bacteria pairs will not only resolve phage activities in regulating hyphosphere life, but also offer a powerful tool for testing hypotheses of invasion ecology at large scale (e.g. regarding spatial and trait relationships of alien biota) in spatially tailored microbial model systems.

## ACKNOWLEDGMENTS

This study is part of the Collaborative Research Centre AquaDiva of the Friedrich Schiller University Jena, funded by the Deutsche Forschungsgemeinschaft (DFG, German Research Foundation) - SFB 1076 - project number 218627073 and the Helmholtz Centre for Environmental Research - UFZ. The authors wish also to thank Maria Fabisch and Anke Hädrich for great coordination of the CRC AquaDiva and iRTG AquaDiva. The authors are thankful for the use of the helium-ion microscope at the Centre for Chemical Microscopy (ProVIS) at UFZ Leipzig, which is supported by European Regional Development Funds (EFRE—Europe funds Saxony) and the Helmholtz Association.

## SUPPLEMENTARY MATERIAL

Supporting Information is available and contains 7 pages with 4 figures.

## CONFLICT OF INTEREST

The authors declare that the research was conducted in the absence of any commercial or financial relationships that could be construed as a potential conflict of interest.

## Notes

### Competing Interest Statement

The authors have declared no competing interest.

