## Supporting information for "Phage co-transport with hyphal-riding bacteria fuels bacterial invasion in water-unsaturated microbial ecosystems"

**Summary**: The supporting information includes 7 pages and 4 figures

**Extended Materials and Methods**

Time-dependent transport rates ($R_i$), transport capacity ($C_i$) and transport efficiency ($E_i$) was used to evaluate phage-bacterial co-transport in microcosm setups shown in Fig. 2a. Eqs. S1 & S2 define the cumulative transport rates of bacteria ($R_b$, CFU cm$^{-1}$ d$^{-1}$) and phages ($R_p$, PFU cm$^{-1}$ d$^{-1}$) at given time (t, d). $N_{b,agarBC}$ (CFU) and $N_{p,agarBC}$ (PFU) reflect the number of infectious phages or bacteria on agar patches B & C, $d$ (cm) the distance between inoculum and edge of the agar strip, and $t$ (d) the time interval between inoculation and sampling. As T4 got rapidly deactivated on agar surfaces (99% loss of PFU in < 24 h, Fig. S2a) and subsequent low phage numbers were elusive to direct quantification, $N_{p,agarBC}$ (eq. S2) was approximated by the difference of phage numbers on agar patch A in the absence ($N_{p,agarA}$) and presence ($N'_{p,agarA}$) of carrier bacteria. This approach was possible as the presence of *P. ultimum* or *P. ultimum* and *P. putida* KT2440 co-cultures did not influence T4 infectivity (Fig. S2c).

$$R_b = \frac{N_{b,agarBC}}{d \times t} \qquad \text{eq. S1}$$

$$R_p = \frac{N_{p,agarBC}}{d \times t} \approx \frac{N_{p,agarA} - N'_{p,agarA}}{d \times t} \qquad \text{eq. S2}$$

The average number of phages co-transported by a single bacterial carrier, i.e. the apparent bacterial phage transport capacity ($C_p$, PFU bacteria$^{-1}$), is reflected by eq. S3.

$$C_p = \frac{N_{p,agarBC}}{N_{b,agarBC}} \approx \frac{N_{p,agarA} - N'_{p,agarA}}{N_{b,agarBC}} \qquad \text{eq. S3}$$

The fraction of phages dispatched by carrier bacteria, i.e. phage transport efficiency ($E_p$, %), was calculated by eq. S4.

$$E_p = \frac{N_{p,agarA} - N'_{p,agarA}}{N_{p,agarA}} \qquad \text{eq. S4}$$

Eq. S5 refers to the time-dependent absolute fitness ($W_i$) [37] in order to reflect the effects of phage co-transport on the fitness of bacterial ($W_b$) or phage ($W_p$) populations in microcosm setups shown in Fig 3a. $N_b$ (CFU) and $N_p$ (PFU) represent the phage (i.e. T4) or bacteria (i.e. *P. putida* KT2440) counts on agar patches A, B or C in the absence of *E. coli*. $N^*_b$ (CFU) and $N^*_p$ (PFU) reflect the phage (i.e. T4; $N_{T4}$) or bacteria (i.e. *P. putida* KT2440; $N_{WT}$) on agar patches A, B or C in presence *E. coli*. $W_i > 1$ and $W_i < 1$ indicate an increase and a decrease, resp. of the population size, while $W_i = 0$ refers to population extinction. At $t = 1$ d, no significant difference (P>0.05, Fig. 2d) between $N_{p,agarA}$ and $N'_{p,agarA}$ (i.e. an estimation for $N_{p,agarBC}$) was observed in the absence of *E.coli*. We thus assigned the PFU detection limit (= 200 PFU $ml^{-1}$) to $N_{p,agarBC}$ for a conservative estimation of $W_p$ on day 1.

$$W_b = \frac{N^*_b}{N_b} \qquad W_p = \frac{N^*_p}{N_p} \qquad \text{eq. S5}$$

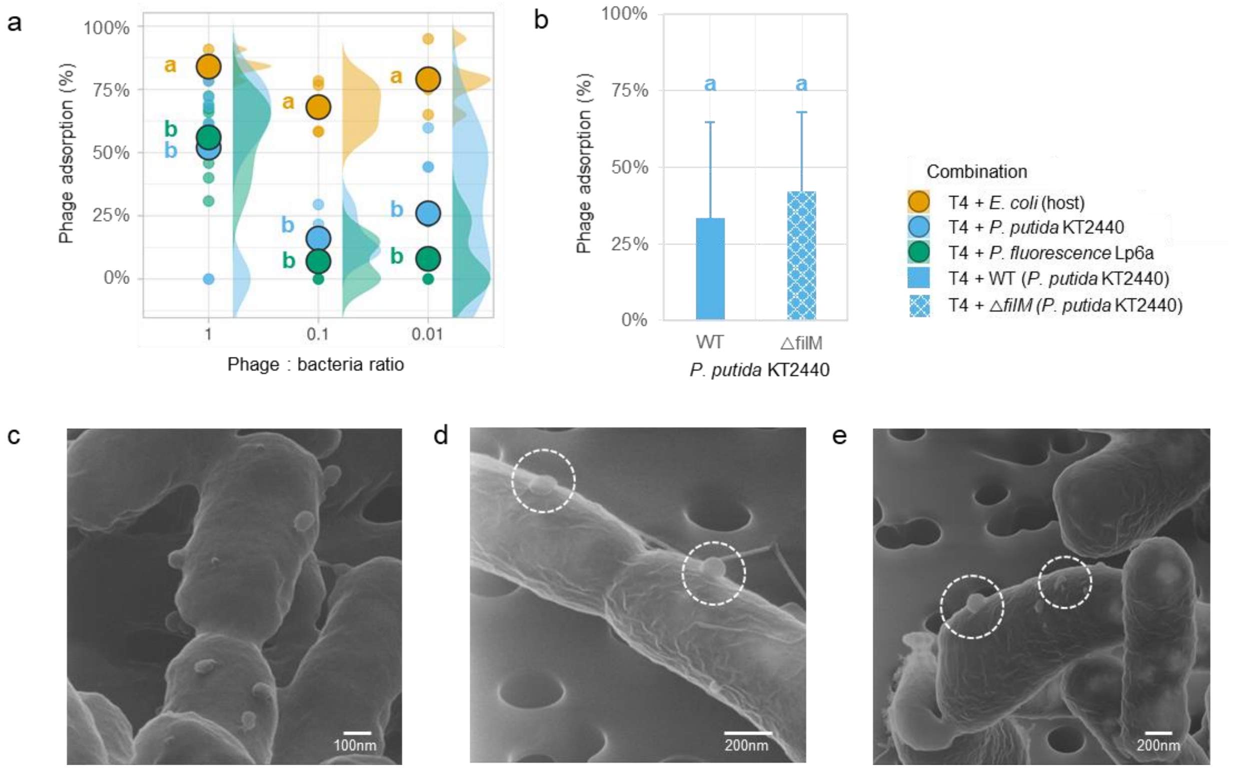


**Figure S1**. T4 adsorption to host *E. coli* bacteria,non-host bacteria *P. putida* KT2440 (WT and non-flagellated △*film*), and *P. fluorescence* Lp6a cells in liquid adsorption assays (cf. materials and methods); **Fig S1a**. Adsorption of T4 to *E. coli* host and non-host WT and *P. fluorescens* Lp6a at 3 tested phage to bacteria ratios (i.e. 0.01, 0.1 and 1). Data represent 6-8 replicates at each phage-to-bacteria ratio and different letters denote statistical significance at P<0.05 (Welch's *t* test); **Fig S1b**. Adsorption of T4 to flagellated WT and non-flagellated △*film* at a phage to bacteria ratio of 1. Different letters denote statistical significance at P<0.05 (n = 3; Welch's *t* test). **Fig S1c**. HIM visualization of T4 adsorption to *E.coli*. **Fig S1d** & **S1e**. HIM visualization of T4 adsorption to WT.

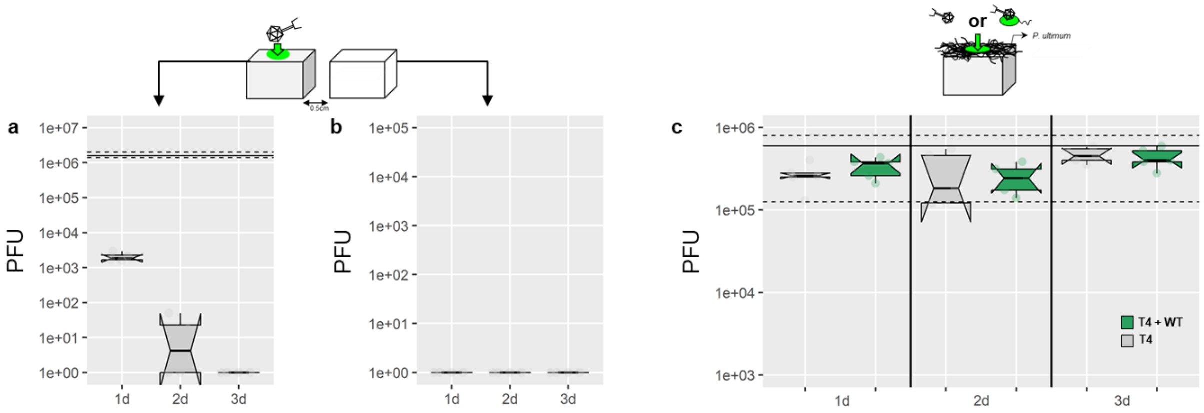


**Figure S2**. Time dependent abundance of infectious T4 on agar surfaces in presence and absence of additional biomass (*P. ultimum* or *P. ultimum* + *P. putida* KT2440). **Fig S2a&b**. Decay of T4 on an agar surface over time (Fig. S2a; bold and dashed lines indicate the median ± 95% confidential interval of the inoculated amount; n = 5) and air-born transport of T4 to another agar surface at 0.5 cm distance (Fig S2b). **Fig S2c**. Time-dependent decay of T4 on an agar surface overgrown either by *P. ultimum* or *P. ultimum* + *P. putida* KT2440 (bold and dash lines indicate the median ± 95% confidential interval of the inoculated amount; n = 5).

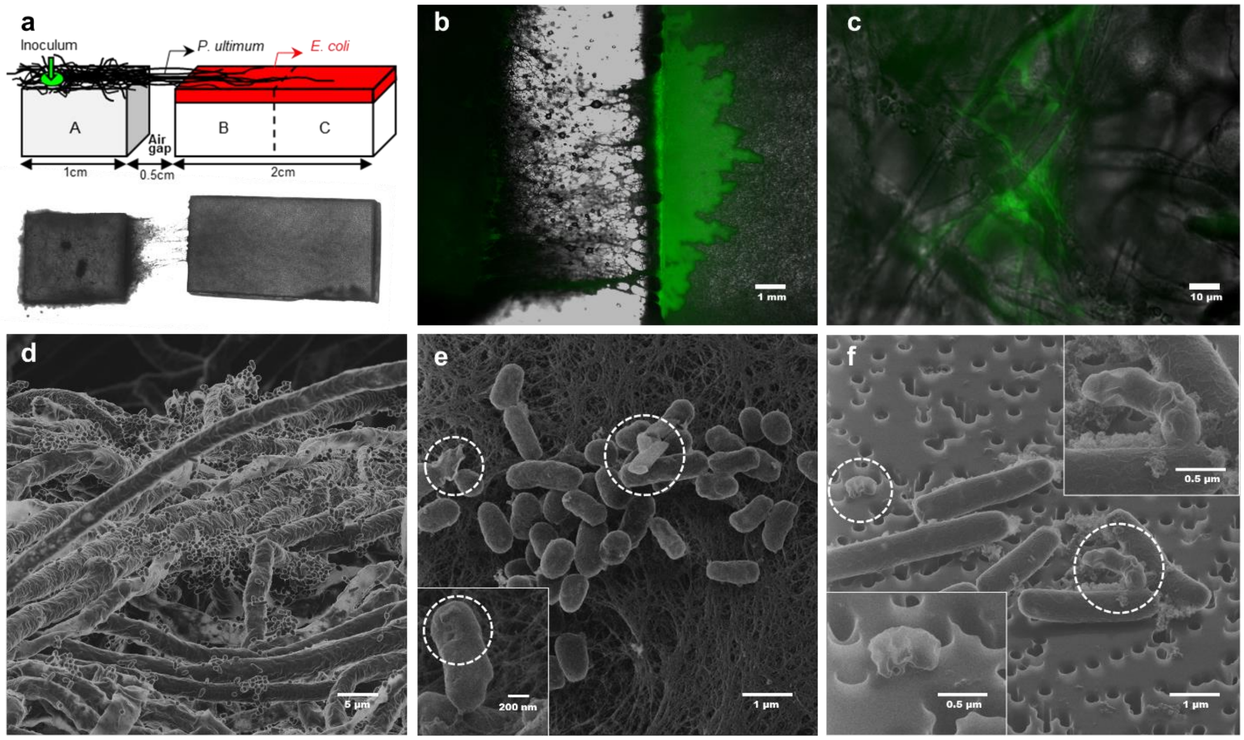


**Figure S3**. Microscopic visualization of T4 adsorbed *P. putida* KT2440 (WT) invading into an *E. coli* biofilm (**Figs S3b&c**), *P. putida* KT2440 (WT) in the hyphosphere of *P. ultimum* (**Figs S3d**), and T4 induced lysis of *E. coli* cells (**Figs S3e&f**). **Fig S3a**. Scheme and top view photograph of the microcosm setup. **Fig S3b**. Representative fluorescence micrograph (60% opacity of gfp overlay) depicting the air-gap between agar patches A & B and invasion of T4 adsorbed and gfp-labelled WT (in green) to *E. coli* biofilm on agar patch B. Fig **S3c**. Representative fluorescence micrograph (60% opacity of gfp overlay) depicting establishment and growth of WT (in green) in the hyphosphere on agar patch C on day 3. **Fig. S3d-f**. Helium ion micrographs visualizing E. coli and WT in the hyphosphere of *P. ultimum* (**Fig. S3d**) and lysed *E. coli* cells on agar patch B on day 2 (**Fig. S3e**) and of controls of T4 infected *E.coli* cells after 25 min of co-incubation with T4 in liquid medium (**Fig. S3e**).

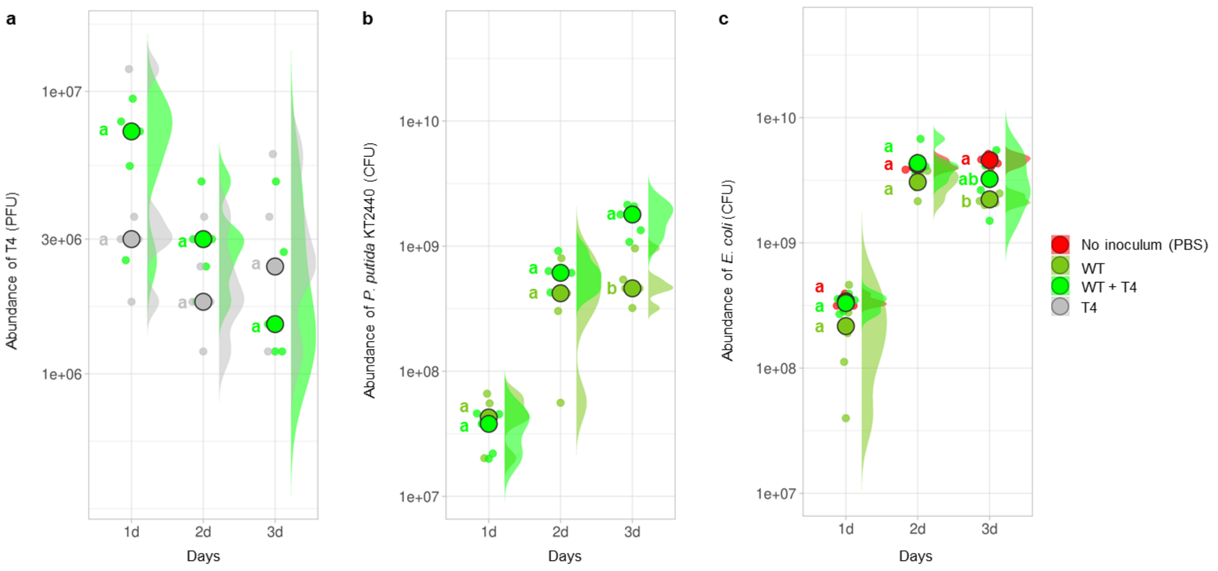


**Figure S4**. Abundances of T4, *P. putida* KT2440 (WT) and *E. coli* in the whole microcosm (agar patches A, B & C). **Fig. S4a**. Abundances of T4 in the presence and absence of the WT carrier. **Fig. S4b**. Abundances of WT in the presence and absence of T4. **Fig. S4c**. Abundances of *E.coli* in the absence of WT and in presence of either WT or WT + T4. Data include 5 replicates and different letters denote statistical significance at $P < 0.05$ (Welch's *t* test).
